# Characterisation of a second gain of function *EDAR* variant, encoding EDAR380R, in East Asia

**DOI:** 10.1101/813063

**Authors:** Jon Riddell, Chandana Basu Mallick, Guy S. Jacobs, Jeffrey J. Schoenebeck, Denis J. Headon

**Author notes:** Centre for Genetic Disorders, Institute of Science, Banaras Hindu University, Varanasi, India (Chandana Basu Mallick).

## Abstract

EDAR is a TNF receptor family member with roles in the development and growth of hair, teeth and glands. A derived allele of *EDAR*, single nucleotide variant rs3827760, encodes EDAR370A, a receptor with more potent signalling effects than the ancestral EDAR370V. This allele of rs3827760 is at very high frequency in modern East Asian and Native American populations as a result of ancient positive selection and has been associated with straighter, thicker hair fibres, alteration of tooth and ear shape, reduced chin protrusion and increased fingertip sweat gland density. Here we report the characterisation of another SNV in *EDAR*, rs146567337, encoding EDAR380R. The derived allele of this SNV is at its highest global frequency, of up to 5%, in populations of southern China, Vietnam, the Philippines, Malaysia and Indonesia. Using haplotype analyses, we find that the rs3827760 and rs146567337 SNVs arose on distinct haplotypes and that rs146567337 does not show the same signs of positive selection as rs3827760. From functional studies in cultured cells, we find that EDAR380R displays increased EDAR signalling output, at a similar level to that of EDAR370A. The existence of a second SNV with partly overlapping geographic distribution, the same *in vitro* functional effect and similar evolutionary age as the derived allele of rs3827760, but of independent origin and not exhibiting the same signs of strong selection, suggests a northern focus of positive selection on EDAR function in East Asia.

## Introduction

Ectodysplasin A1 receptor (EDAR) is a cell surface receptor involved in the development of ectodermal structures including hair, teeth and glands (Headon and Overbeek, 1999). Upon activation by its ligand Ectodysplasin (EDA), EDAR signals through its cytoplasmic adapter protein EDARADD to trigger the activation of the transcription factor NF-ĸB (Headon et al., 2001), this signalling sequence being essential for its developmental function (Schmidt-Ullrich *et al.*, 2001). Disruption of this highly conserved signalling pathway via loss-of-function mutations of any of *EDA*, *EDAR* or *EDARADD* causes hypohidrotic ectodermal dysplasia (HED) (Kere *et al.*, 1996; Monreal *et al.*, 1999; Headon *et al.*, 2001), a condition characterised by sparseness of hair, the loss or reduction of many skin-associated glands, and tooth agenesis (Reyes-Reali *et al.*, 2018). Selective absence of teeth commonly occurs as a result of milder function-reducing mutations in the EDA-EDAR pathway, without eliciting the complete set of clinical HED phenotypes (Arte *et al.*, 2013).

The death domain of the EDAR protein is essential for the recruitment of the EDARADD protein and thus for EDAR function (Headon *et al.*, 2001). This domain is present in many proteins, the majority of which are involved in cell death and inflammation (Park *et al.*, 2007). The death domain is approximately 80 amino acids in length and is composed of six alpha helices, these forming a surface that is capable of self-association and of binding to other specific death domain containing proteins (Park *et al.*, 2007; Ferrao and Wu, 2012).

A non-synonymous single nucleotide variant (SNV), rs3827760 (*EDAR:*c.1109T>C), encodes a valine to alanine substitution within the death domain of EDAR at amino acid position 370. The derived allele is at very high frequency in northern East Asian and Native American populations, with allele frequencies of up to 90% in some groups (The 1000 Genomes Project Consortium, 2015). The *EDAR:*c.1109T>C allele displays clear evidence of positive selection both from haplotype and allele frequency spectrum based analyses (Carlson *et al.*, 2005; Sabeti *et al.*, 2007; Bryk *et al.*, 2008; Myles *et al.*, 2008; Grossman *et al.*, 2010; Cheng, Xu and DeGiorgio, 2017); at least some of this selection presumably occurred in the common ancestors of modern East Asian and Native American populations. EDAR370A has been shown to increase the activation of NF-ĸB compared to that of the protein encoded by the ancestral allele (EDAR370V) *in vitro* using reporter assays (Bryk et al., 2008; Mou et al. 2008), and ameliorate the clinical signs of HED caused by hypomorphic *EDA* mutations in heterozygous carriers of EDAR370A (Cluzeau *et al.*, 2012), strongly indicating that the derived allele is a gain-of-function. The physiological consequences of this increased signalling have been assessed in mouse models, either with multiple copies of *EDAR* to increase expression level and signalling, or through engineering of the *EDAR:*c.1109T>C variant in mice (Mou *et al.* 2008; Kamberov *et al.*, 2013). Both of these models were observed to have thicker hair fibres and, complementing these findings, human association studies have shown that *EDAR:*c.1109T>C is associated with thicker, straighter scalp hair, along with other traits such as shovelling of incisors, altered ear and chin shape, and increased fingertip sweat gland density (Fujimoto *et al.*, 2008; Kimura *et al.*, 2009; Kamberov *et al.*, 2013; Tan *et al.*, 2013; Adhikari *et al.*, 2015; Adhikari, Fontanil, *et al.*, 2016; Adhikari, Fuentes-Guajardo, *et al.*, 2016; Wu *et al.*, 2016; Shaffer *et al.*, 2017).

Here we identify another SNV in *EDAR* (rs146567337, *EDAR*:c.1138A>C) which causes a serine to arginine substitution at amino acid position 380 (EDAR380R). The geographic distribution of the derived allele of this SNV partly overlaps that of the previously characterised *EDAR:*c.1109T>C (encoding EDAR370A), though at lower frequency and with a more southerly prevalence. The EDAR380R substitution increases the signalling function of EDAR to a similar degree as the EDAR370A substitution, but its genomic context does not show the same signs of strong positive selection in human populations, despite both alleles having approximately the same age (Albers and McVean, 2019). These findings suggest that *EDAR*:c.1138A>C (EDAR380R) may influence the same human traits as those associated with *EDAR:*c.1109T>C (EDAR370A), and that these traits may have been under different selective pressures in different regions of Asia.

## Results

The death domain of EDAR is a highly conserved region of the protein, with mutations altering this domain commonly leading to a loss-of-function and thus clinically diagnosed hypohidrotic ectodermal dysplasia (Cluzeau *et al.*, 2011), presumably due to altered or abrogated EDAR interaction with EDARADD (Okita *et al.*, 2019). We identified SNV rs146567337 (*EDAR*:c.1138A>C) in *EDAR* in the gnomAD database (https://gnomad.broadinstitute.org/) (Karczewski *et al.*, 2019). The derived allele encodes a serine to arginine substitution at the highly conserved amino acid 380 (EDAR380R), only 10 amino acids from the alteration in the well-characterised EDAR370A variant (Fig 1A). In the gnomAD database (Karczewski *et al.*, 2019), which is not a representative sampling of worldwide populations, the frequency of the derived allele at rs146567337 was 1.85%. Using publicly available datasets (The 1000 Genomes Project Consortium, 2015; Mallick *et al.*, 2016; Pagani *et al.*, 2016; Jacobs *et al.*, 2019), we found *EDAR*:c.1138A>C only in East and Southeast Asian populations, at highest frequency in southern China, Vietnam, the Philippines, Malaysia, and Indonesia. However, the distribution of this allele did not extend further south and east into New Guinean populations (Fig 1B and 1C). Since *EDAR:*c.1109T>C is at very high frequency in many of the populations with appreciable frequencies of *EDAR*:c.1138A>C, we assessed whether *EDAR*:c.1138A>C and *EDAR:*c.1109T>C appear on the same haplotype. Using the same datasets, we analysed haplotypes spanning a 20 kb window surrounding rs146567337, on which *EDAR*:c.1138A>C is present and found only one occurrence, out of 33 assessed *EDAR*:c.1138A>C haplotypes from 5,608 evaluated chromosomes, where *EDAR:*c.1109T>C and *EDAR*:c.1138A>C co-existed on the same haplotype. This singular occurrence, possibly a genotyping or phasing error, suggests that the derived allele of rs146567337 arose on a different haplotype to that of *EDAR:*c.1109T>C (Fig 1D). We also found in modern human populations the entire haplotype context of the *EDAR*:c.1138A>C allele, but with the ancestral allele at this SNV, likely representing the immediately ancestral haplotype to *EDAR*:c.1138A>C on which this mutation occurred. This immediately ancestral haplotype is geographically widely dispersed, in East Asian, South Asian, African, and American populations. The variant was also found to be ancestral in both the Altai Neanderthal (Prüfer *et al.*, 2014) and Altai Denisovan (Meyer *et al.*, 2012) genomes, and was not inferred to be in archaic introgressed haplotypes identified in either the Simons Genome Diversity Project (Mallick *et al.*, 2016) or Indonesian Genome Diversity Project data (Jacobs *et al.*, 2019). Hence, we infer that the *EDAR*:c.1138A>C variant arose in modern humans rather than through introgression from an archaic human population.

**Figure 1:**
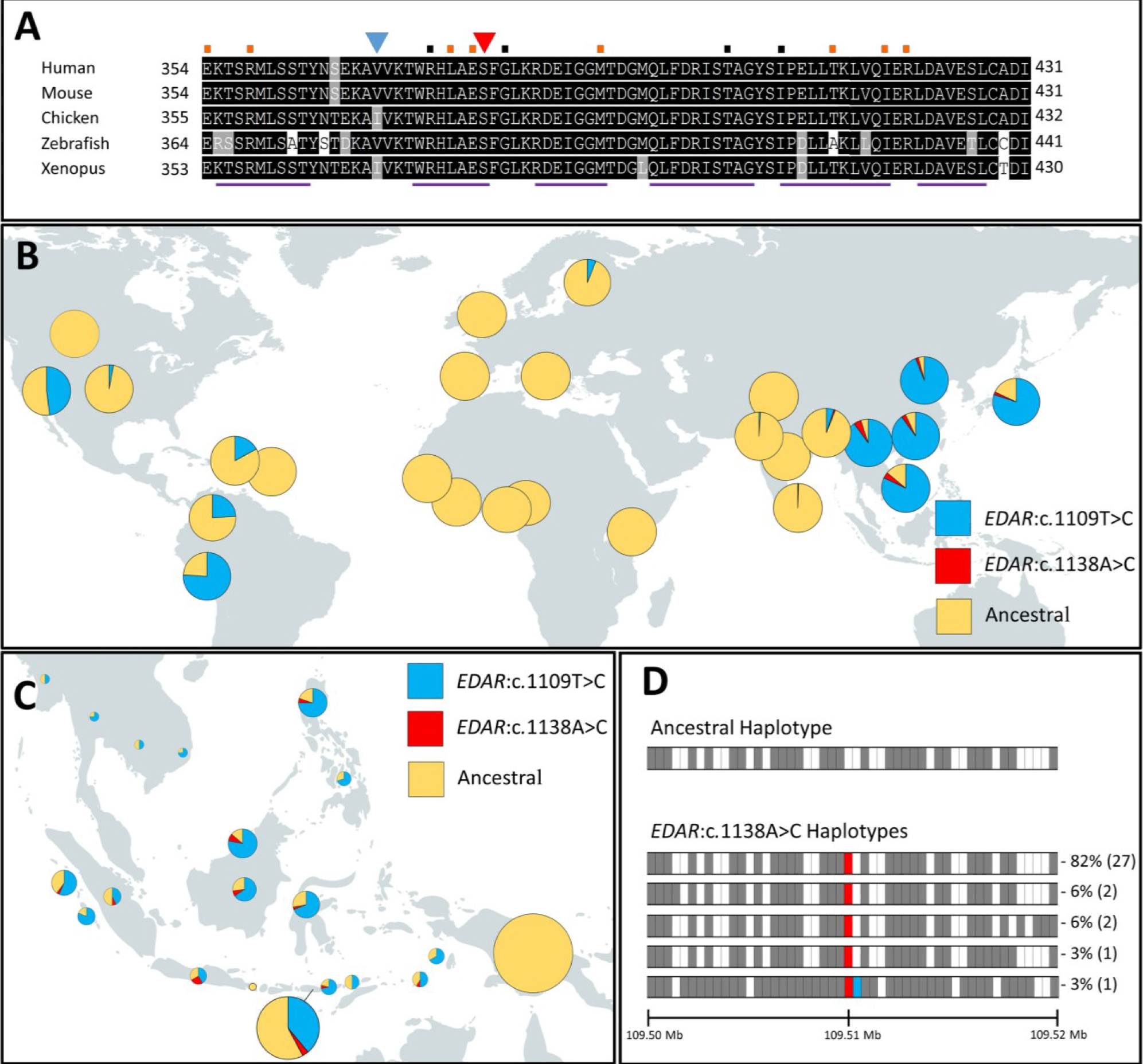
Conservation, distribution and haplotype structure of *EDAR* variants. **(A)** Multiple sequence alignment of vertebrate EDAR death domains. Amino acid positions within the EDAR protein are numbered at the start and end of the sequence for each species. The position of 370 is indicated by a blue triangle, the position of 380 by a red triangle. The positions of known recessive and dominant mutations causing hypohidrotic ectodermal dysplasia in humans are indicated by black and orange squares, respectively, above the alignment. Purple bars below the alignment indicate the positions of the predicted alpha helices. **(B)** Worldwide allele frequencies for *EDAR:*c.1109T>C and *EDAR*:c.1138A>C in the 1000 Genomes dataset plotted as pie charts for each population. The remaining allele frequency was depicted as ancestral. **(C)** *EDAR:*c.1109T>C and *EDAR*:c.1138A>C allele frequencies in the Southeast Asian Island populations were gathered from publicly available datasets (Pagani *et al.*, 2016; Jacobs *et al.*, 2019) and plotted on a map of Southeast Asia as in **(B)**. The area of each chart is proportional to the sample number of each population. **(D)** Diagram of the *EDAR*:c.1138A>C haplotypes from the 1000 Genomes Project and Simons Genome Diversity Project datasets plotted for a region 10 kb upstream and 10 kb downstream of rs146567337. White boxes indicate alleles matching the reference human genome (GRCh37) and grey boxes indicate the presence of the alternate allele. Red shading indicates the *EDAR* 1138C allele and blue shading indicates *EDAR* 1109C. In total, 33 *EDAR*:c.1138A>C haplotypes were present in these datasets, with five unique haplotype structures identified. These were ranked in order of frequency, as shown by the percentages to the right of each haplotype, with the total number of each individual haplotype in the dataset indicated in brackets. The scale bar indicates position on chromosome 2.

To further define the distribution of *EDAR* haplotypes, we constructed a median-joining haplotype network consisting of 142 SNPs spanning about ±10 kb around SNVs of interest (rs3827760 and rs146567337) using a publicly available dataset (Pagani *et al.*, 2016) (Fig 2). We used the HapMap combined genetic map (Frazer *et al.*, 2007) to confirm that the window did not have especially fast recombination likely to disrupt the network reconstruction (0.0797cM; 44^th^ percentile of total genetic map distance in non-overlapping genome wide 114.9kb windows). The network identified 88 haplotypes and further supports the independent origins of *EDAR*:c.1138A>C and *EDAR*:c.1109T>C. The haplotype associated with *EDAR*:c.1109T>C is mainly composed of individuals from East Asia, Siberia, Southeast Asia Island and mainland populations, and the Americas (Fig 2). Individuals from Siberia (denoted by cyan) represent almost 50% of this haplotype in this population sample. The dataset used for the construction of this network sampled few East Asian individuals (n=11) than Siberians (n=108), thus explaining the greater proportion of the latter with the associated haplotype (Pagani *et al.*, 2016). We also observed that the *EDAR*:c.1109T>C associated haplotype demonstrates a star-like pattern, suggestive of a historic demographic expansion and corroborating earlier evidence of positive selection at this locus. In contrast, the haplotype associated with *EDAR*:c.1138A>C was found to be distant from *EDAR*:c.1109T>C and showed more restricted geographic distribution, confined to individuals mainly from the islands of Southeast Asia and one individual from South Asia.

**Figure 2:**
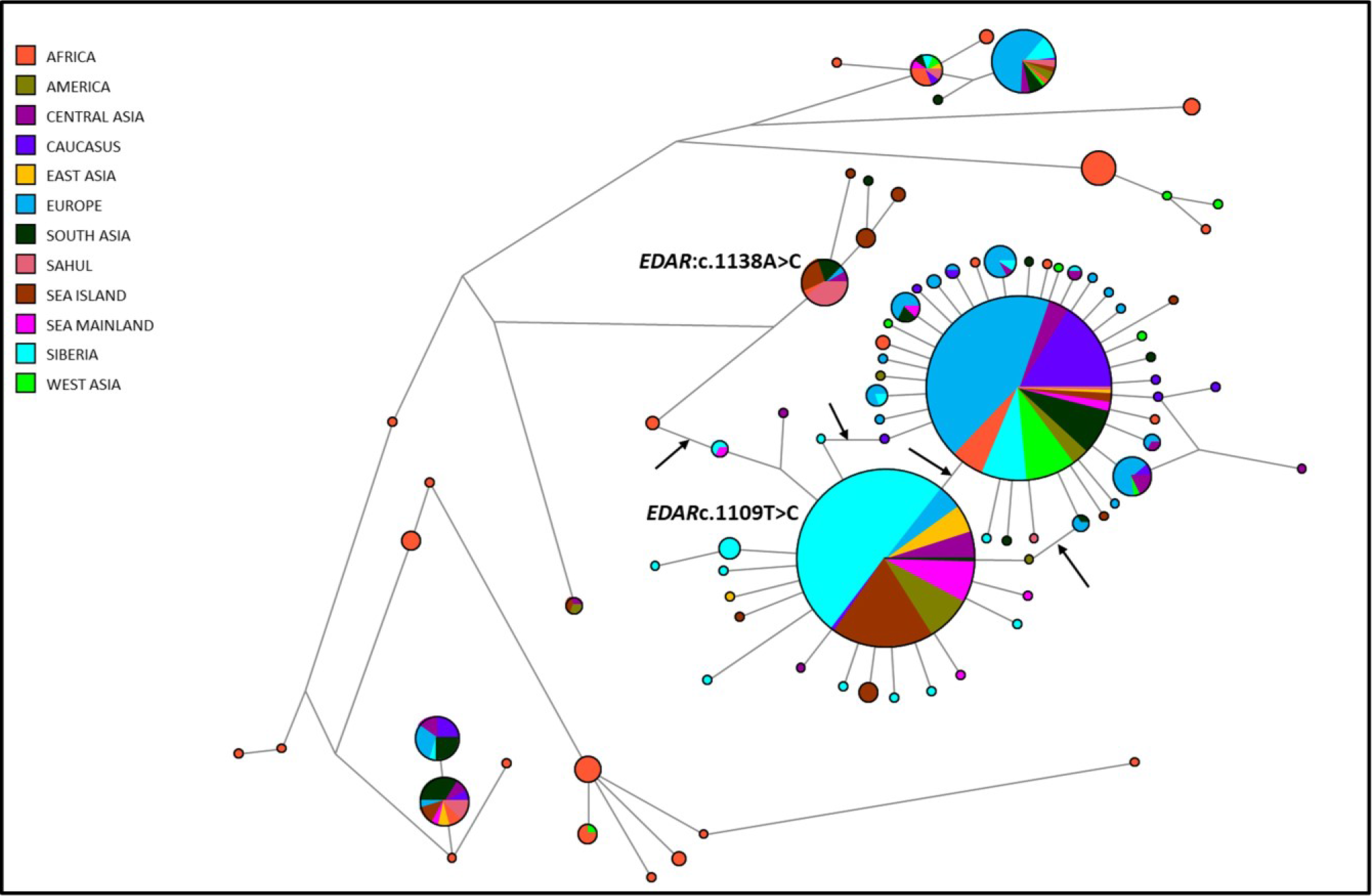
Median-joining haplotype network of *EDAR*. Median-joining haplotype network spanning ± 10 kb around rs3827760 and rs146567337 showing the relationship of the haplotypes. The network is based on 446 individuals included in the Pagani *et al.*, 2016 dataset. Each pie chart represents a unique haplotype and the size of the chart is proportional to the number of chromosomes carrying it. Colours represent the geographic location of populations where each haplotype was found. Lines represent mutations, with greater branch length indicating a greater number of distinguishing mutations. The associated haplotypes of interest (for *EDAR:*c.1109T>C and *EDAR*:c.1138A>C) have been labelled. The black arrows represent locations where an *EDAR:*c.1109T>C mutational event was inferred. The multiple arrows likely reflect ambiguity in the network reconstruction.

As *EDAR:*c.1109T>C is one of the most well-supported examples of a positively selected locus in the human genome (Carlson *et al.*, 2005; Kelley *et al.*, 2006; Sabeti *et al.*, 2007; Xue *et al.*, 2009; Hider *et al.*, 2013), we tested for indications of selection on *EDAR*:c.1138A>C. Using a large-scale whole genome sequence dataset of the Han Chinese population (Cai *et al.*, 2017), we constructed extended haplotype homozygosity (EHH) plots of 433 *EDAR*:c.1138A>C haplotypes (derived rs146567337, ancestral rs3827760) and 20,293 *EDAR:*c.1109T>C haplotypes (derived rs3827760, ancestral rs146567337) against 554 double ancestral haplotypes (haplotypes bearing the ancestral alleles for both rs146567337 and rs3827760) (Fig 3A). No double derived allele haplotypes were found in this Han Chinese dataset. As expected for loci that underwent selection, and as demonstrated previously (Sabeti *et al.*, 2002, 2007), *EDAR:*c.1109T>C shows a broad region of haplotype homozygosity compared to the double ancestral haplotype. *EDAR*:c.1138A>C exhibits much less extended haplotype homozygosity than *EDAR:*c.1109T>C, despite being at a lower derived allele frequency, suggesting that *EDAR*:c.1138A>C has not been subjected to the same pressures or degree of selection as *EDAR:*c.1109T>C. The Han Chinese dataset included 433 *EDAR*:c.1138A>C (EDAR380R) haplotypes, therefore we constructed EHH bifurcation plots by random subsampling of 433 haplotypes from double ancestral allele and *EDAR:*c.1109T>C (EDAR370A) haplotypes. The bifurcation plots confirmed that the *EDAR*:c.1138A>C haplotype had been reduced by recombination less frequently than the double ancestral haplotype, but more frequently than the *EDAR:*c.1109T>C haplotype (Fig 3B).

**Figure 3:**
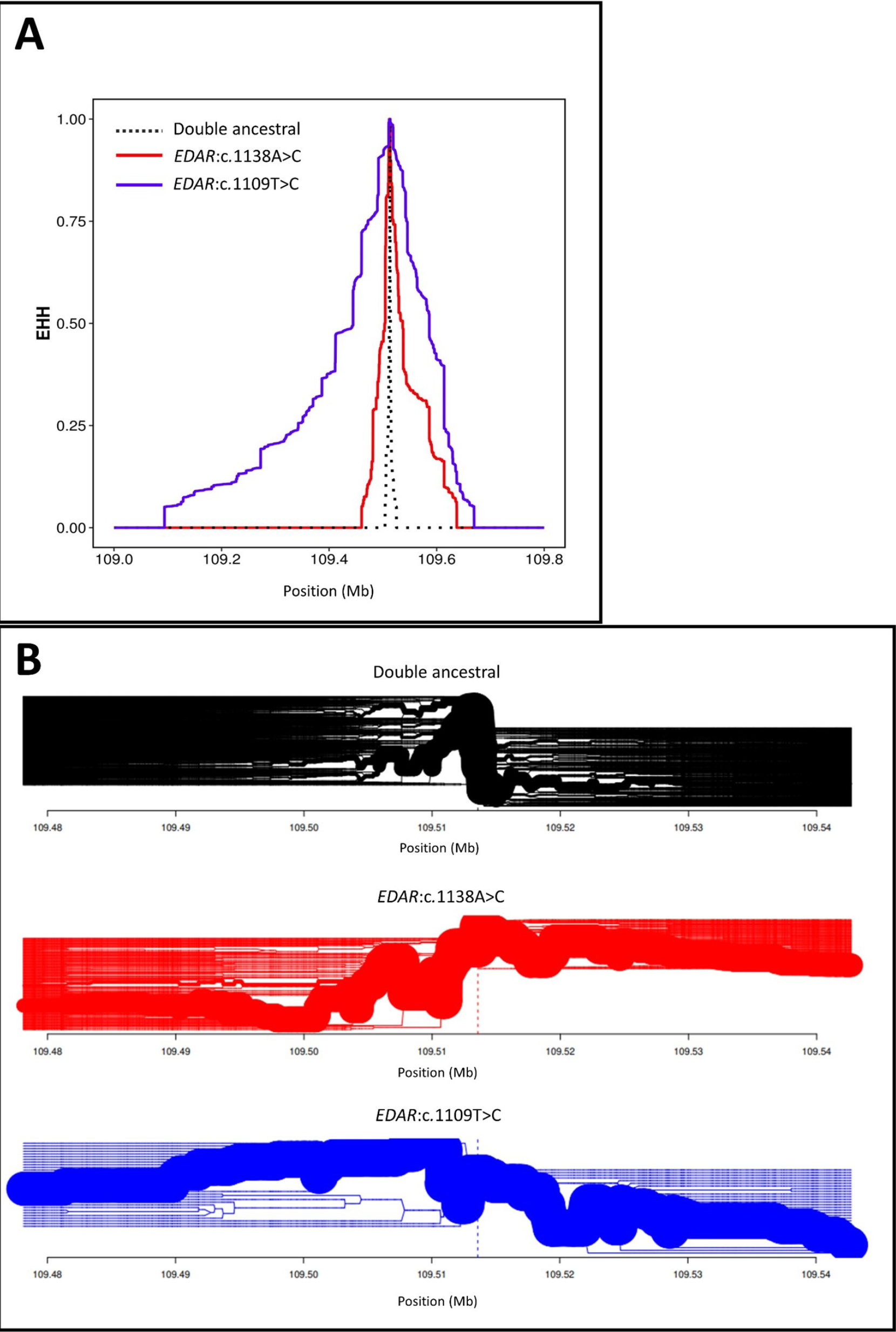
Extended haplotype homozygosity (EHH) and EHH bifurcation plots surrounding *EDAR* variants. EHH plot showing the length of conserved haplotype on either side of rs146567337. An EHH value of 1 indicates that haplotypes are identical at this position. Double ancestral haplotypes are represented by the black dotted line, *EDAR*:c.1138A>C (EDAR380R) haplotypes are represented by the red line, and *EDAR:*c.1109T>C (EDAR370A) haplotypes are represented by the blue line. **(B)** Bifurcation plot showing the branching of each haplotype. Thicker lines indicate more common haplotypes. Double ancestral haplotypes are represented by the black line, *EDAR*:c.1138A>C haplotypes are represented by the red line, and *EDAR:*c.1109T>C haplotypes are represented by the blue line.

After determining the global distribution and genomic context of *EDAR*:c.1138A>C, we next investigated the effect of the substitution on the encoded protein. To map the position of EDAR380R within the death domain and identify any predicted structural effects of this amino acid substitution, we modelled the variant EDAR death domain structures using the intensive mode of the Phyre2 server. This program uses multiple homologous templates to predict the structure of a given input protein sequence (Kelley *et al.*, 2015). The resulting predicted protein structure positioned amino acid EDAR380 within an alpha helix (Fig 4A), a structural feature known to be important for the protein-protein interactions mediated by death domains (Ferrao and Wu, 2012). However, the alternate amino acid variants did not alter the predicted structure of this helix or any other part of the death domain. The protein structure also remained unaltered when we modelled the EDAR370A substitution (Fig 4A). Taken together, based on the conservation of the serine residue at position EDAR380 among vertebrates (Fig 1A), introduction of a positive charge through its substitution to arginine and strong evidence of functional alteration reflected from SIFT (score 0) (Ng and Henikoff, 2003) and PolyPhen (score 0.999) (Adzhubei *et al.*, 2010), we predicted that the EDAR380R substitution would alter EDAR protein function. To test this, we transfected HEK293T cells, a human cell line derived from embryonic kidney, with *EDAR* cDNAs encoding either the ancestral EDAR, EDAR370A, EDAR380R, or the double substituted EDAR370A+EDAR380R protein. We also included the known loss-of-function variant EDAR379K, which is dominant for selective tooth agenesis in humans and autosomal recessive for HED in mice (Headon and Overbeek, 1999; Arte *et al.*, 2013), as a control in these experiments. Each form was assayed for its ability to activate a co-transfected NF-ĸB luciferase reporter. We found that EDAR380R activated NF-ĸB in these cells to a greater degree than ancestral EDAR, and to the same extent as EDAR370A, and that the greatest activation of NF-ĸB was observed when EDAR carried both the 370A and 380R amino acid substitutions (Fig 4C). These effects on signalling activity were broadly confirmed in the human HaCaT cell line, derived from the skin’s epidermis (Fig 4D).

**Figure 4:**
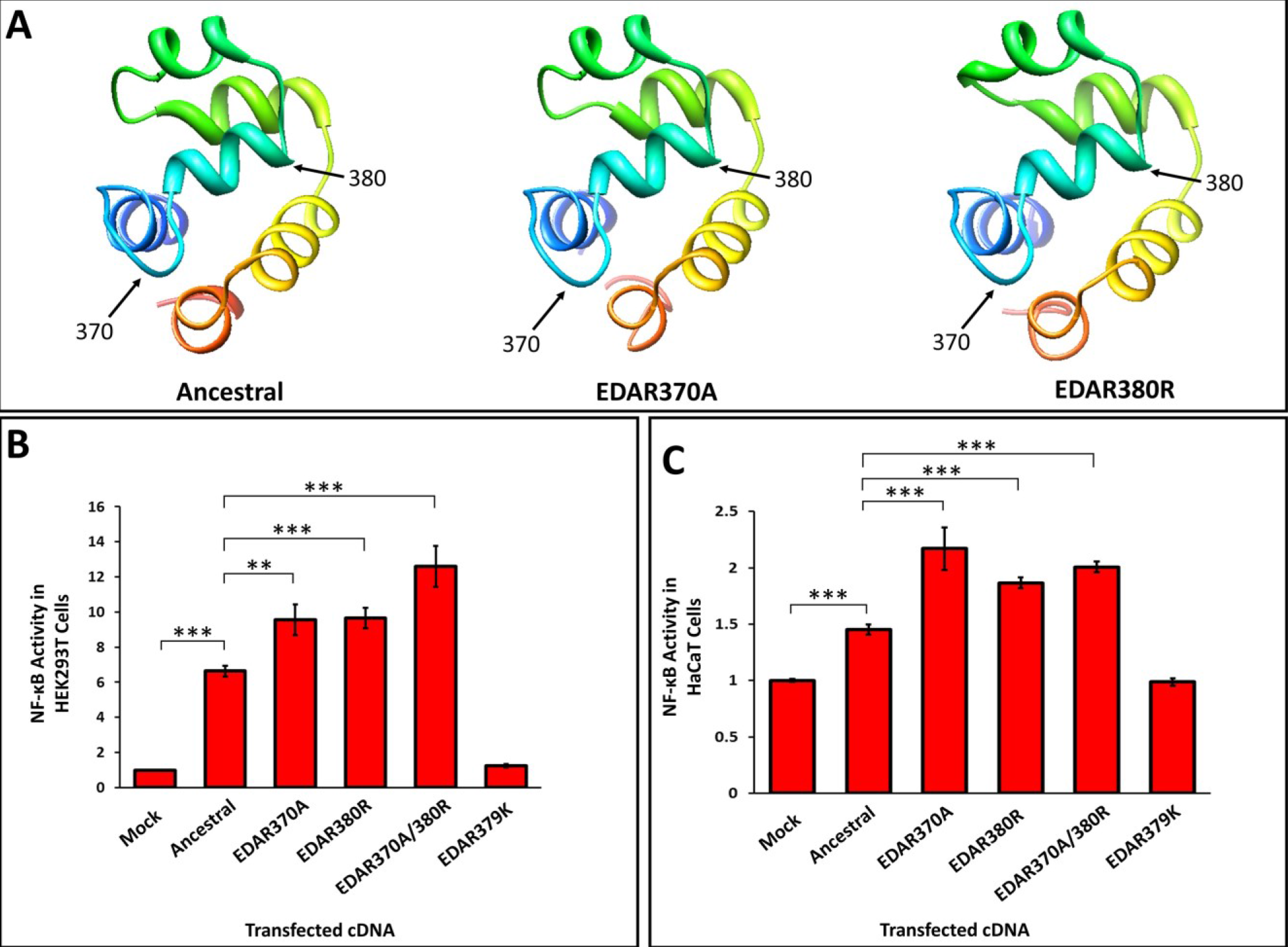
Functional effects of *EDAR* variants. **(A)** The predicted protein structures of the ancestral, EDAR370A and EDAR380R death domains, modelled using Phyre2. The location of amino acid positions 370 and 380 are indicated by arrows. Amino acid position 380 is located towards the end of an alpha helix. **(C)** HEK293T and **(D)** HaCaT cells were transfected to express *EDAR* variants and resulting NF-ĸB luciferase reporter activity determined. Error bars represent the standard error of the mean from experiments performed in quadruplicate and repeated independently 6 times. Statistical significance was calculated using a Student *t* test (** P < 0.005, *** P <0.0005).

## Discussion

We identified and characterised a novel functional variant in *EDAR* through haplotype analyses and cell-based experiments. We find that *EDAR*:c.1138A>C has its highest allele frequency in Southeast Asia and appears to have arisen on a different haplotype to that of the more common and previously characterised *EDAR:*c.1109T>C substitution. We find that *EDAR*:c.1138A>C does not show the same signs of having been under strong positive selection as *EDAR:*c.1109T>C based on extended haplotype homozygosity analyses, but that the encoded protein increases NF-ĸB activation *in vitro*, to approximately the same extent as the *EDAR:*c.1109T>C substitution.

Several theories as to what the selective advantage conferred by EDAR370A was have been advanced. Chang et al. suggested that EDAR370A was positively selected in the ancestors of East Asians and Native Americans for adaptation to a cold and dry climate, in which increased skin-associated glands and resulting glandular secretions, perhaps together with straighter hair, could be advantageous in producing a functional barrier to the environment (Chang *et al.*, 2009). Hlusko et al. suggested a latitude-based adaptive scenario, in which altered transfer of nutrients, particularly vitamin D, through breast milk in far northeast Asia (Hlusko *et al.*, 2018) was caused by the mammary gland alterations enacted by enhanced EDAR signalling (Chang *et al.*, 2009; Kamberov *et al.*, 2013). Kamberov et al. placed the origin of the EDAR370A encoding allele in central China at greater than 30,000 years ago, and suggested that increased eccrine sweat gland number, associated with the *EDAR:*c.1109T>C variant in mouse and human in their study, as the potential selective force that would have been advantageous in hot and humid climates due to increased ability to perspire (Kamberov *et al.*, 2013).

A recent genealogical estimation of allele ages in humans assessed the derived alleles *EDAR:*c.1109T>C and *EDAR*:c.1138A>C as having a very similar date of origin, at approximately 1,400 generations ago (Albers and McVean, 2019). The geographic distribution of these alleles is somewhat similar, and, though at much lower frequency than *EDAR:*c.1109T>C in all regions, it is notable that the highest frequencies of *EDAR*:c.1138A>C overlap the more southerly regions in which *EDAR:*c.1109T>C is prevalent. The *EDAR*:c.1138A>C variant is notably absent from the Americas, where in native populations *EDAR:*c.1109T>C is essentially at fixation (Bryk et al., 2008). We found that *EDAR*:c.1138A>C does not show as strong a signal of positive selection in human populations as *EDAR:*c.1109T>C, despite cell culture experiments predicting similar outcomes resulting from the EDAR370A and EDAR380R substitutions. The frequency of the *EDAR*:c.1138A>C variant peaks in Southeast Asia and it is thus most likely to have arisen in that region. The *EDAR:*c.1109T>C variant appears to have arisen further north, based on its present-day population distribution and ancient DNA analyses (Mathieson *et al.*, 2015; Siska *et al.*, 2017; Lamnidis *et al.*, 2018), suggesting that phenotypes associated with an EDAR-dependent increase in NF-ĸB activation have been preferentially selected for in more northern regions of Asia. The overlap between these alleles could be influenced by frequency dependent selection, perhaps on a phenotype directly perceptible by others.

The possibility that EDAR370A and EDAR380R may have similar phenotypic effects should be considered in future gene or genome-wide association studies, particularly in populations in which the derived allele of rs3827760 is at high frequency. In these populations, only a small fraction of ancestral rs3827760 alleles exist and a sizeable proportion of these haplotype will carry the derived rs146567337 allele, which could obscure the phenotypic associations with the derived allele of rs3827760. The *EDAR*:c.1138A>C allele has been identified in people exhibiting tooth agenesis in East Asian populations (He *et al.*, 2013; Yamaguchi *et al.*, 2017). However, our data suggest that this allele is unlikely to be causative for the condition due to its increased, rather than decreased, activity, as observed for *EDAR*:c.1109T>C.

The discovery of a second SNV in *EDAR* that increases NF-ĸB activation to the same extent as *EDAR*:c.1109T>C raises questions as to how many routes there are to achieving the same molecular effect of increased EDAR activity. Multiple mutations have also been identified in an enhancer region upstream of the *LCT* gene that have the same molecular effect of increasing *LCT* transcription (Tishkoff *et al.*, 2007). However, these *LCT* SNVs exhibited clear EHH, suggesting that this molecular effect was selected for in each of the populations containing these variants. In this case, *EDAR*:c.1138A>C does not show the same extent of EHH as *EDAR*:c.1109T>C, even though the alleles are predicted to be of a similar age, and should therefore both have had the opportunity to be selected. This indicates that either the molecular consequences of *EDAR*:c.1109T>C are more complex *in vivo* than reflected in cell signalling assays, or that the phenotypic consequences of enhanced *EDAR* signalling have only been strongly selected for in northern East Asian populations. This work highlights that exploring the population genetics of variants with similar molecular phenotypes as known selected variants could prove beneficial in the future for refining the features of those variants, and the relevant environments, that led to their selection.

## Materials and Methods

### Generation of phylogeographic maps

Maps of the world and of Southeast Asia were generated using MapChart (https://mapchart.net/). The rs3827760 and rs146567337 allele frequencies were gathered from publicly available datasets (The 1000 Genomes Project Consortium, 2015; Pagani *et al.*, 2016; Jacobs *et al.*, 2019) and plotted on to pie charts for each population. The pie charts were used to annotate the maps based on the co-ordinates that samples originated from.

### Determination of archaic human genotypes

We used the high coverage Altai Neanderthal (Prufer et al. 2014; downloaded from http://cdna.eva.mpg.de/neandertal/altai/AltaiNeandertal/VCF/) and Altai Denisovan (Meyer et al. 2012 et al; downloaded from http://cdna.eva.mpg.de/denisova/VCF/hg19_1000g/) to determine the state of rs146567337 in archaic hominins. As the archaic genomes are sampled from populations that are divergent from introgressing archaic populations, we additionally confirmed a non-archaic origin using introgressing archaic haplotypes inferred by Jacobs *et al*. (2019) for samples in the Simons Genome Diversity Project and Indonesian Genome Diversity Project.

### Haplotype analysis

VCF files from publicly available datasets (The 1000 Genomes Project Consortium, 2015; Mallick *et al.*, 2016) were narrowed down to a 20 kb window surrounding rs146567337. These files were viewed using inPHAP software (v1.1) (Jäger, Peltzer and Nieselt, 2014) and the variants that disagreed with the human reference genome (GRCh37) were mapped for samples containing *EDAR*:c.1138A>C. This process was repeated for samples containing *EDAR:*c.1109T>C and the list of variants mapped were combined to give a total of 50 SNVs that were used for the final haplotype constructs.

### Construction of the median-joining haplotype network of *EDAR*

A median-joining haplotype network was constructed using NETWORK 5.0 (http://www.fluxus-engineering.com) to study the phylogenetic relationship between the haplotypes. For this, we used the region around ± 10 kb of the SNVs of interest (rs3827760 and rs146567337) among the individuals included in the Pagani et al. 2016 dataset.

### Generation of extended haplotype homozygosity (EHH) and bifurcation plots

EHH plots and bifurcation plots were constructed from data generated in a large scale whole-genome sequencing dataset of 11,670 individuals from the Han Chinese population (Cai *et al.*, 2017) using R package rehh (v2.0.2) (Sabeti *et al.*, 2002). The EHH plot was generated by separating the haplotypes into three categories: Ancestral - where the ancestral alleles of both rs3827760 and rs146567337 are present, *EDAR*:c.1138A>C - where the derived allele of rs146567337 and the ancestral allele of rs3827760 are present, *EDAR:*c.1109T>C - where the ancestral allele of rs146567337 and the derived allele of rs3827760 are present. In total there were 554 double ancestral haplotypes, 433 *EDAR*:c.1138A>C haplotypes and 20,293 *EDAR:*c.1109T>C haplotypes. In order to aid visualisation in the bifurcation plot, 433 haplotypes from each category were randomly selected and plotted.

### Sequence alignment and protein structure modelling

The EDAR death domain sequence alignment was generated with the T-coffee alignment tool (Notredame, Higgins and Heringa, 2000; Di Tommaso *et al.*, 2011) using peptide sequences gathered from the NCBI protein database (GenInfo Identifiers: Human - 11641231, Mouse - 6753714, Chicken - 60302666, Zebrafish - 924859488, Xenopus - 55742031). The generated fasta_aln file was then entered into BOXSHADE v3.21 to obtain a shaded output indicating amino acid sequence conservation.

The EDAR death domain protein structure was generated using the intensive mode of Phyre2 v2.0 (Kelley *et al.*, 2015) by inputting amino acid positions 345-431 of the human EDAR peptide sequence gathered from the NCBI protein database (GenInfo Identifier: 11641231).

### Transfection of cells and luciferase assays

HEK293T and HaCaT cells were maintained at 37°C in 5% CO_2_ in high glucose Dulbecco’s modified Eagle’s medium (DMEM) supplemented with 10% foetal bovine serum (FBS) and 50 μg/ml streptomycin and 100 U/ml penicillin (Gibco). Transfections of HEK293T and HaCaT cells were performed using Lipofectamine 3000 (Invitrogen) in 24-well plates (well surface area: 1.9 cm^2^). Both cell lines were seeded at a density of 5 × 10_4_ 24 hours prior to transfection. Each well was transfected with a DNA mix diluted in opti-MEM (Gibco) consisting of 125 ng pNFĸB-luc, 62.5 ng pRLTK, 10 ng pCR3::EDAR, pCR3::EDAR370A, pCR3::EDAR380R, pCR3::EDAR370A/EDAR380R or pCR3::EDAR379K, and made up to a total amount of 500 ng with empty pCR3.1 vector. Transfections were performed according to manufacturer’s instructions in DMEM supplemented with 10% FBS, 50 μg/ml streptomycin and 100 U/ml penicillin.

Luciferase assays were performed 18 hours post-transfection using the Dual-Luciferase Assay System (Promega) according to manufacturer’s instructions, using a MicroLumatPlus LB96V Microplate Luminometer.

